# A novel decoding strategy for ProteinMPNN to design with less MHC Class I immune-visibility

**DOI:** 10.1101/2025.04.14.648837

**Authors:** Hans-Christof Gasser, Ajitha Rajan, Javier A. Alfaro

## Abstract

Due to their versatility and diverse production methods, proteins have attracted a lot of interest for industrial as well as therapeutic applications. Designing new therapeutics requires careful consideration of immune responses, particularly the cytotoxic T-lymphocyte (CTL) reaction to intra-cellular proteins. In this study, we introduce `CAPE-Beam`, a novel decoding strategy for the established `ProteinMPNN` protein design model. Our approach minimizes CTL immunogenicity risk by limiting designs to only consist of kmers that are either predicted not to be presented to CTLs or are subject to central tolerance. We compare `CAPE-Beam` to greedily sampling from `ProteinMPNN` and `CAPE-MPNN`. We find that our novel decoding strategy can produce structurally similar proteins while incorporating more human like kmers. This significantly lowers CTL immunogenicity risk in precision medicine, and represents a key step towards reducing this risk in protein therapeutics targeting a wider patient population.

## 1 Introduction

Fueled by recent advances in AI methods, the field of computational protein design has experienced much growth and attention recently. Protein design foundation models like `ProteinMPNN` [6] are being used to predict a protein’s amino acid (AA) sequences based on its desired 3D structure. As of today, rarely any of these models take the reaction of the immune-system to these designed sequences into account. This is not a problem for proteins designed for industrial applications. However, in case new therapeutics should be developed using these methods, this will have to change. In particular, as AI-designed intracellular protein therapeutics incorporate novel functions and diverge from native sequences, they may be increasingly recognized by CTLs via the MHC Class I (MHC-I) pathway, potentially reducing efficacy or triggering autoimmune responses, as seen with anti-transgene immunity [4, 18].

There are two sequence-based approaches that we can think of to reduce the immune-reaction by CTLs to a therapeutic protein. CTLs detect their targets based on short kmers (mostly 8-10mers [20, Section 4-15]) that make up the protein’s AA sequence and are presented to them on the cell’s surface (see **Subsection 2.1**). In addition, to avoid auto-immunity, CTLs are trained to ignore peptides that are parts of human proteins. Therefore, kmers that are not presented on the cell surface to CTLs and kmers that are part of the human proteome (self kmers) should not result in an immune-reaction by CTL.

Based on this, the first approach utilizes the property of the MHC-I system, not to present all peptides on the cell surface. By avoiding the incorporation of presented peptides into protein designs, we can potentially hide the therapeutic from the person’s CTLs. Based on *netMHCpan 4*.*1* [21] predictions with a rank cutoff of 2%, we estimate in this work that c10% of the 8 to 10-mers would be presented in a single individual. However, this number increases significantly to cover a significant part of the population. Designing complex protein structures without a large proportion of all potential kmers, seems to be difficult. Therefore, we believe that this approach will only be workable for precision medicine solutions that focus on a single patient.

Interestingly, while the 8-10mers in the human proteome only cover a tiny slice of all possible kmers (less than 0.02%) — they still give rise to the full complexity of human biology. This motivates the hypothesis that a wide range of structures can be constructed using only self kmers. CTLs should then be tolerant to all peptides resulting from the degradation of these proteins. An advantage is that – except for some variation in the human genome – the resulting drugs do not have to be personalized. In general, making the protein sequence look more similar to human proteins seems to reduce immune-reactions as highlighted by the fact that T-cell receptors (TCRs) bind far less strongly to tumor-associated antigenic peptides, which have sequences very similar to those found in the human proteome, than to viral ones [5]. Also, Richman et al. found that cancer neoantigen dissimilarity to the self-proteome predicts immunogenicity [22].

Hence, we introduce `CAPE-Beam`. This novel decoding strategy for `ProteinMPNN` aims to either incorporate kmers from the human genome (self kmers) or not presented kmers into the generated designs. We found that restricting only to self kmers resulted in very low predicted TM-scores for the generated designs. However, when only restricting to self 6-mers while longer kmers were allowed to be non-human (non-self kmers) but not presented, then high predicted TM-scores can be achieved.

## 2 Background

We now briefly discuss the detection of foreign intra-cellular proteins by the adaptive immune-system as well as two key machine learning (ML) elements for this paper. The first is `ProteinMPNN` – the foundation model used by our decoding strategy. The second are some basic decoding strategies for auto-regressive (AR) models like `ProteinMPNN`.

### 2.1 Detection of intra-cellular proteins [20]

Proteins have a wide range of functions in biology. They are able to fulfill those, due to their flexible, diverse design. This is based on chains of AAs that, due to the different biochemical properties of these AAs, fold into different sequence dependent 3D structures in solutions. Within a cell, proteins are constantly being produced and degraded – leaving short AA sub-chains behind. The MHC-I pathway presents a subset of these (mostly about 8-10 AAs long) on the cell surface. Which ones will depend on the MHC-I alleles expressed by the individual. These presented peptides are then subject to surveillance by CTLs. These are trained in the thymus to tolerate peptides present in the human proteome - thereby preventing auto-immunity. In contrast, foreign proteins can cause destruction by CTL under the right conditions.

### 2.2 ProteinMPNN

`ProteinMPNN` [6] is a protein design model that takes as input a desired 3D protein-backbone structure (the template) and, in a AR fashion, generates a corresponding AA sequence in response. With its roughly 1.7m parameters it is a comparatively small model. It also supports the design of complexes consisting of multiple AA chains.

`ProteinMPNN` consists of two message passing neural networks (MPNNs) - one *Encoder* and one *Decoder*. The *Encoder* first encodes the available information into a representation that can then be used by the *Decoder* to generate the sequence iteratively. The order in which the sequence is decoded can be specified arbitrarily, allowing for fixing certain parts of the sequence upfront. This paper examines an alternative approach to sample from the AA distribution generated by the network in each step.

### 2.3 Decoding strategies

An AR language model typically outputs the logit probability values for each of the potential next categories/tokens to sample from. The traditional way to choose the next token is then to either always select the most likely next token (greedy strategy - the “standard” strategy in this paper), or to construct a Boltzmann distribution (softmax) with a temperature parameter (using the negative logits as energy levels) to randomly sample the next token from. Variants of this are *top-k* sampling, in which only the most probable *k* next tokens would be considered for sampling. Or, *top-p* sampling, which only considers the smallest set of tokens that has a cumulative probability of more than *p*.

A very different strategy is *beam search*. The origins of which can be traced back at least to the 1970s and the first implementation is often attributed to [16]. Here, *k* sequence generations are considered in parallel and then all the following possible continuations are expanded. So if we had 21 possible next tokens, then there would be 21 *× k* candidate continuations. From these, only the *k* most likely ones are kept and the process is repeated. Based on this idea, we formulated the `CAPE-Beam` decoding strategy. For the interested reader, a list of decoding strategies and a comparison of them with regards to large language models (LLMs) can be found in Shi et al. [23].

## 3 Related Work

The exact problem solved by computational protein design is ill defined. Often times, the inverse folding problem is meant - finding a AA sequence that folds into a given 3D structure. However, sometimes people also just refer to the generation of a AA sequence based on other constraints as computational protein design.

With regards to the inverse folding problem, traditional physics based approaches were searching for AA sequences that would minimize a scoring function for a given confirmation. The score is typically at least partly modeling the potential energy of a confirmation. However, it would regularly also include other terms - like statistical ones that could account for evolutionary relationships. The *Rosetta Packer* [15, 8] is one of these traditional physics-based approaches using Monte-Carlo simulation to search for optimal AA sequences. Yachnin et al. [27] modified the Rosetta score function by adding an additional term to reduce visibility to helper thymus dependent lymphocytes (T-cells) via the MHC Class II (MHC-II) pathway. This approach was taken up and modified in the `CAPE-Packer` [10] to modify visibility to CTLs via the MHC-I pathway.

The recent advances in ML approaches have reinvigorated the computational protein design process. Instead of minimizing a score function, these methods learn the sequence-structure relationship from data. Often a diffusion model like *RFdiffusion* [25] or FoldingDiff [26] would generate a 3D protein structure. The inverse-folding problem of finding a corresponding AA sequence to this is then handled by models like `ProteinMPNN` [6] or *ESM-IF* [11].

Most work that used or adapted these methods to consider immune-reactions, has been targeted towards reducing antibody (Ab) reaction. For example Lyu et al. [17] have used a Variational Autoencoder (VAE) approach to generate new adenovirus capsid proteins to prevent vector detection by pre-existing Abs. Similarly, Bootwala et al. [2] have used the framework set out in [29] and the structure-based protein design graph neural network (GNN) model developed by Ingraham et al. [12] to re-surface the L-Asparaginase protein to avoid Ab detection.

In contrast, our recent work introduced `CAPE-XVAE` (based on a VAE architecture) [10] and `CAPE-MPNN` [9] (fine-tuning `ProteinMPNN` [6]), to specifically reduce visibility to CTLs. However, none of this work was explicitly trying to incorporate kmers from the human proteome to leverage central tolerance. This paper, therefore, introduces a novel decoding strategy that explicitly prefers sampling self kmers.

## 4 Methods

This section first sets out the way we assess presentation of peptides by the MHC-I pathway. Then we discuss the `CAPE-Beam` decoding strategy before presenting the ways we evaluated the `CAPE-Beam` designed sequences.

### 4.1 Predicting presented peptides

One of the most commonly used bioinformatics tools for assessing presentation of peptides on the cell surface via MHC-I is *netMHCpan 4*.*1* [21]. It takes a peptide and a MHC-I allele as input and indicates the likelihood of cell surface presentation via a score. Based on this a rank is calculated which compares the peptide’s score to the one of random natural peptides. A rank below 2% includes “weak binders”, while the stricter threshold of 0.5% corresponds to “strong binders”. [21]

As calculating *netMHCpan* predictions can be a bit slow, we constructed an approximate, fast position weight matrix (PWM) based classifier for which more details can be found in [9]. We use this only during sequence design, while *netMHCpan* is still directly used to assess the presentation of the 8-10mers during evaluation. PWMs are commonly used in bioinformatics. In contrast to [9] we are here generating a PWM for each MHC-I allele and peptide length (8-10) separately.

### 4.2 CAPE-Beam **decoding strategy**

Our method is a novel decoding strategy for `ProteinMPNN`, which ensures that kmers in the generated sequence are either part of the human proteome - in which case central tolerance should prevent an immune-reaction by CTLs - or that they are not presented to CTLs at all.

Inspired by the classic beam search (see **Subsection 2.3**), we generate `width` number of sequence designs in parallel (see **Figure 1**). In accordance with the standard way used in `ProteinMPNN,` we first sample the unresolved (missing coordinates) positions of the input template structure. Afterwards, we sample the remaining positions in an increasing order.

**Figure 1.**
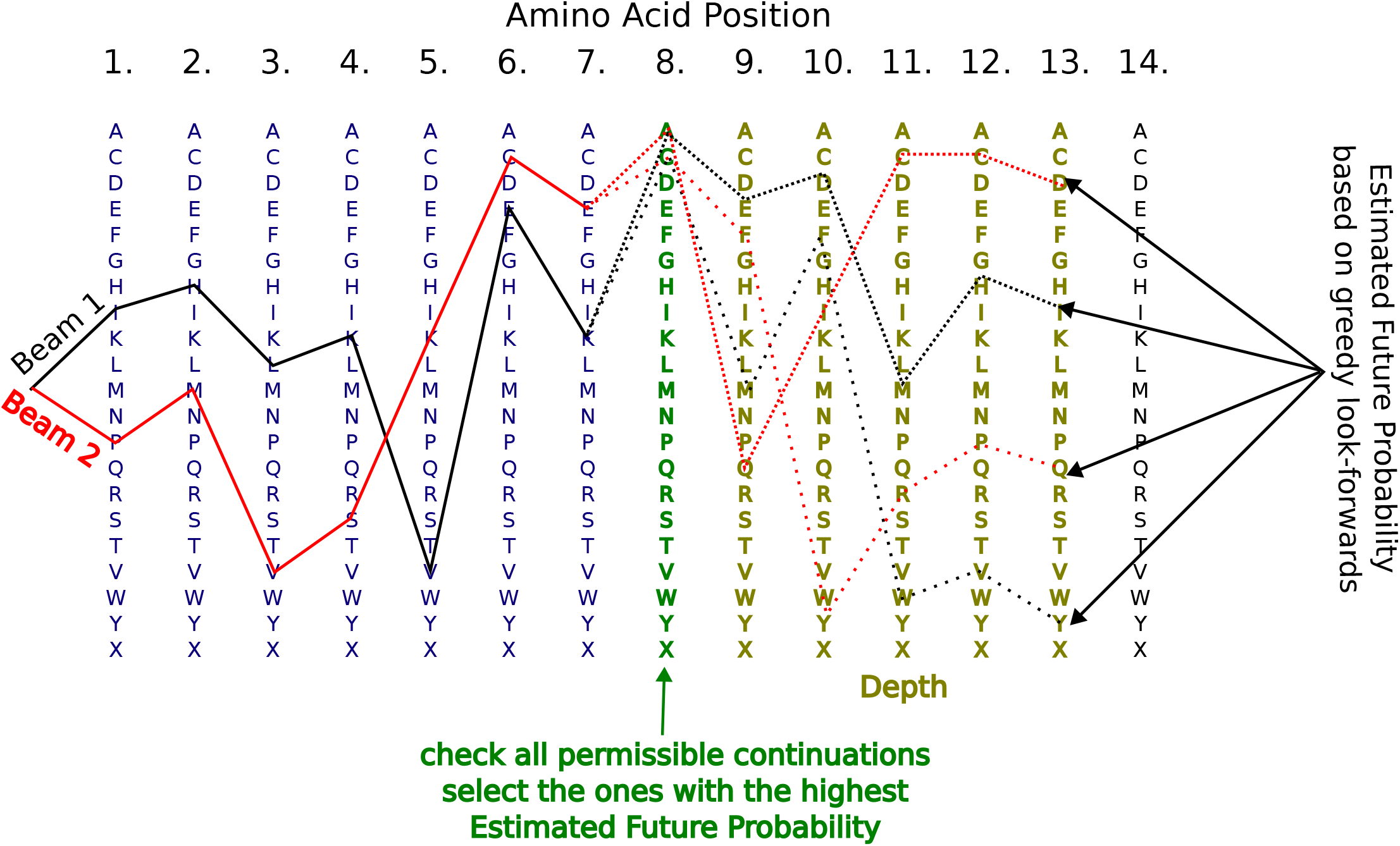
Protein sampling. AAs are added to the protein sequence one after the other. At every newly sampled position (green) all possible extensions are checked for permissibility. For all permissible extensions, we sample a single permissible forward looking sequence continuation (in dashed lines) for up to `n_depth` steps. For every beam (Beam 1 in black and Beam 2 in red), there will be up to 21 of those. The width ones with the highest estimated future probability are kept and expanded further until all residues are determined.

To extend a beam by one token, there are 21 (all standard AA plus ‘unknown’) candidate extensions. To assess the permissibility of an extension, we check that all newly emerging kmers in it (up to a length of `max_checked_kmer_length`) are permissible. Let us assume we had a `max_checked_kmer_length` of 4. If for example we had already sampled IHLKVEK in the beam and were now considering adding the candidate token A, then we would check A, KA, EKA and VEKA (see **Figure 1** Beam 1). There are two ways for a kmer to be permissible. Up to a length of `min_self_kmer_length`, a kmer needs to come from the human proteome (be part of any human protein - we used GRCh38, Ensembl v94). Kmers of length in between `min_self_kmer_length` and `max_checked_kmer_length` can either be self kmers or be predicted not to be presented by the MHC-I pathway (see **Subsection 4.1**). If the extension leads to a non-self kmer (but not presented), then its estimated *future probability* (see below) is multiplied by a penalizing factor (`non_self_prob_factor` times `depth`). This is supposed to favor continuations using self kmers instead of relying on predicted non presentation. There can be a maximum of `width` times 21 remaining permissible beam extensions. The width ones with the highest *future probability* constitute the new beams which are then extended again until all residues are determined. The beam with the highest likelihood is then used as the design.

The *future probability* referred to above is estimated by continuing a candidate extension for `depth` steps and aggregating the probabilities of sampling the most likely permissible AA at each step (see **Figure 1** ‘estimated future probability’). We only perform a greedy exploration of the most likely permissible continuations and do not penalize non-proteome kmers continuations for this purpose.

To prevent collapsing beams (that the beams differ in the first few positions but then have the same continuations), we group all sequences that have the same last 5 AAs, only keeping the most likely one.

### 4.3 Dataset

We use the six *specific validation proteins* (PDB IDs: 1B9K, 1OA4, 1QWK, 1TJE, 1XGD, 4RQG) from [9] as templates. These were randomly selected from the original `ProteinMPNN` validation set, to ensure that they were not used during `ProteinMPNN` training.

### 4.4 Evaluation

We assessed `CAPE-Beam` designs with regards to three criteria, comparing them with three benchmarks: the original template protein, the standard way of decoding `ProteinMPNN` designs (using a temperature of 10^*−*8^) and CAPE-MPNN designed proteins (using the design method described in [9], checkpoint `458340e4:epoch_20`). The first criteria is, how structurally similar the designs are to the original templates. The next one is, how many kmers in the sequences are presented to the immune-system and not part of the human proteome. Finally, we compare some molecular properties.

#### 4.4.1 Structural similarity

To assess the structural similarlity of our designs to the input protein templates, we first used colabfold, which is based on AlphaFold 2, to predict 3D structures for our generated AA sequences [13, 7, 19]. We use the TM-score as a metric to compare these predicted structures to the templates [28]. Scores below 0.2 are associated with random, unrelated proteins. Scores above 0.5 point to the same fold in SCOP/CATH ^1^

#### 4.4.2 Potentially immunogenic kmers

We evaluate each 8-10mer segment of the AA sequence to determine if it belongs to the human proteome and if it is predicted to be presented on the cell surface.

We use netMHCpan 4.1 [21] with a rank cutoff of 2% to check whether a 8-10mer was presented by any of the up to six MHC-I alleles in a human cell (in contrast to the decoding strategy, which uses a faster approximate method - see **Subsection 4.1**). We then present the number of non-self kmers that are presented by any allele, divided by the total number of 8-10mers. Theoretically, our decoding strategy should guarantee that this number is zero for `CAPE-Beam` designs. However, since we use different predictors during sampling and evaluation, a small difference can emerge.

Inspired by the work of Richman et al. [22] we also examine the similarity of non-self 8-10mers incorporated into the sequences to the human proteome. For these non-self kmers we calculate a “BLOSUM62 dissimilarity”. We do so by first identifying the kmer in the human proteome that has the highest BLOSUM62 score to the non-self kmer (without indels). Then we subtract this number from the BLOSUM62 score of the non-self kmer with itself.

#### 4.4.3 Molecular properties

To predict some of the designs’ molecular properties, we used DESTRESS [24]. It analyzes designs using state of the art (SOTA) predictors for various kinds of molecular properties. We focus our attention on the following ones:

- *Rosetta total energy per AA*: A common assumption in protein design is that a protein will spend most of its time in low energy regions of its conformational space. The well known *Rosetta* suite of tools offers an energy function that combines physical as well as statistical terms (see [3]). Therefore it uses Rosetta Energy Units (REUs) instead of standard energy units and there is no direct conversion to kcal/mol possible. To foster comparisons, we normalize them by the number of AAs in the protein. A typical score for a refined structure lies in between −3 and −1 REU per AA [1, 3].
- *aggrescan3d max:* Aggregation of proteins is a commonly faced issue which is often caused by the exposure of hydrophobic regions on the protein’s surface. An aggregation score is calculated for each residue by aggrescan3d [14]. Lower values mean less risk of aggregation.
- *Isoelectric point:* A higher *H*+ concentration (lower pH) will lead to more of them binding to the protein, resulting in a higher charge. The isoelectric point (pI) is then the pH of the solution at which the protein has neutral charge - at higher pH it will be negatively charged and at lower positively.

## 5 Results

In this section we explore the ability of the `CAPE-Beam` decoding strategy described in **Subsection 4.2** to design proteins with reduced visibility to CTL while maintaining structural fidelity. The visibility to CTL is reduced by making them more similar to human proteins and reducing their presentation via the MHC Class I system. This analysis is based on a hypothetical patient with HLA-A*02:01, HLA-A*24:02, HLA-B*07:02, HLA-B*39:01, HLA-C*07:01 and HLA-C*16:01 MHC-I alleles and carried out on the template proteins described in **Subsection 4.3**.

In **Figure 2** we compare the designs generated using various CAPE-Beam decoding strategy hyper-parameters to the benchmarks (see **Section 4.4**), with regards to structural similarity and immune-reaction risk. The first column of **Figure 2** shows that heavily penalizing not presented non-self kmers (using a very low `non_self_prob_factor`, **CAPE-Beam 0.0**), results in designs that are predicted to look very different from the original templates (low TM scores). However, some freedom to chose these not presented kmers quickly leads to more structurally similar designs. Still, we find that forcing all 7mers or higher to come from the human proteome leads to suboptimal designs – presenting a challenge to de-immunization with respect to 8-10mer peptides in the restricted sequence space of the human proteome. However, we can find many high quality designs when forcing the incorporation of only 6mers that come from the human proteome, which certainly would increase the mimicry to the human proteome. **Figure 2** column two shows that all three benchmarks (template, standard, `CAPE-MPNN`) only significantly incorporated 5mers into their designs. With regards to predicted presented non-self kmers, these should be deterministically zero for all `CAPE-Beam` designs. However, the need to use rapid algorithms to approximate antigen presentation for design compared to evaluation (see **Section 4**), means that this is not the case. In fact, using the same presentation predictor for evaluation as during decoding, does result in no predicted to be presented non-self kmers being incorporated by `CAPE-Beam`. Future work utilizing rapid optimized MHC-I presentation predictors will allow for using the same predictors during evaluation and design.

**Figure 2.**
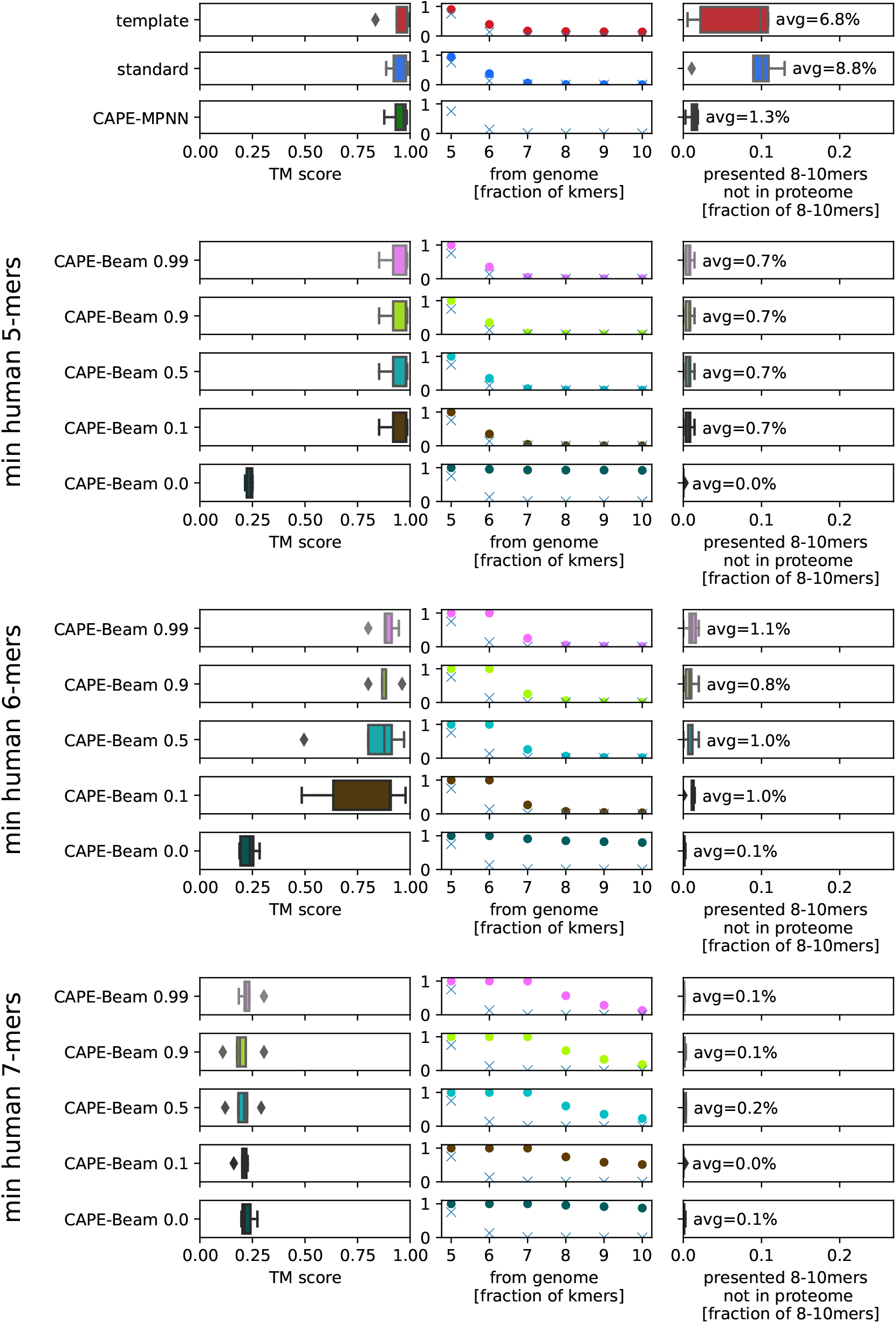
Structural similarity and immune-reaction risk. The first row block depicts the values for the input templates (structure also predicted with AlphaFold to foster like for like comparisons), the designs produced by ProteinMPNN with greedy decoding (“standard”) and the designs produced by `CAPE-MPNN` ([9], 458340e4:epoch 20). Each of the following row blocks is for a different `min_self_kmer_length` parameter. Within these results for `CAPE-Beam` designs using various `non_self_prob_factor` parameters (disclosed after the name) can be found. The first column depicts the distribution of TM scores vs the template protein. The second one shows the percentage of self-kmers (the crosses depict if kmers were uniformly sampled form the standard AAs). The third column then shows the number of presented non-self 8-10mers (potentially immunogenic) as a fraction of all 8-10mers in the protein.

That similarity to the human proteome is relevant from an immunogenicity point of view has been shown in Richman et al. [22]. **Figure 3**, therefore, explores the hypothesis that 8-10mers are more similar to the human proteome in `CAPE-Beam` designs. We find that this is in general the case in comparison to all the benchmarks - except for the template itself, if the protein in question is from the human proteome or a homolog.

**Figure 3.**
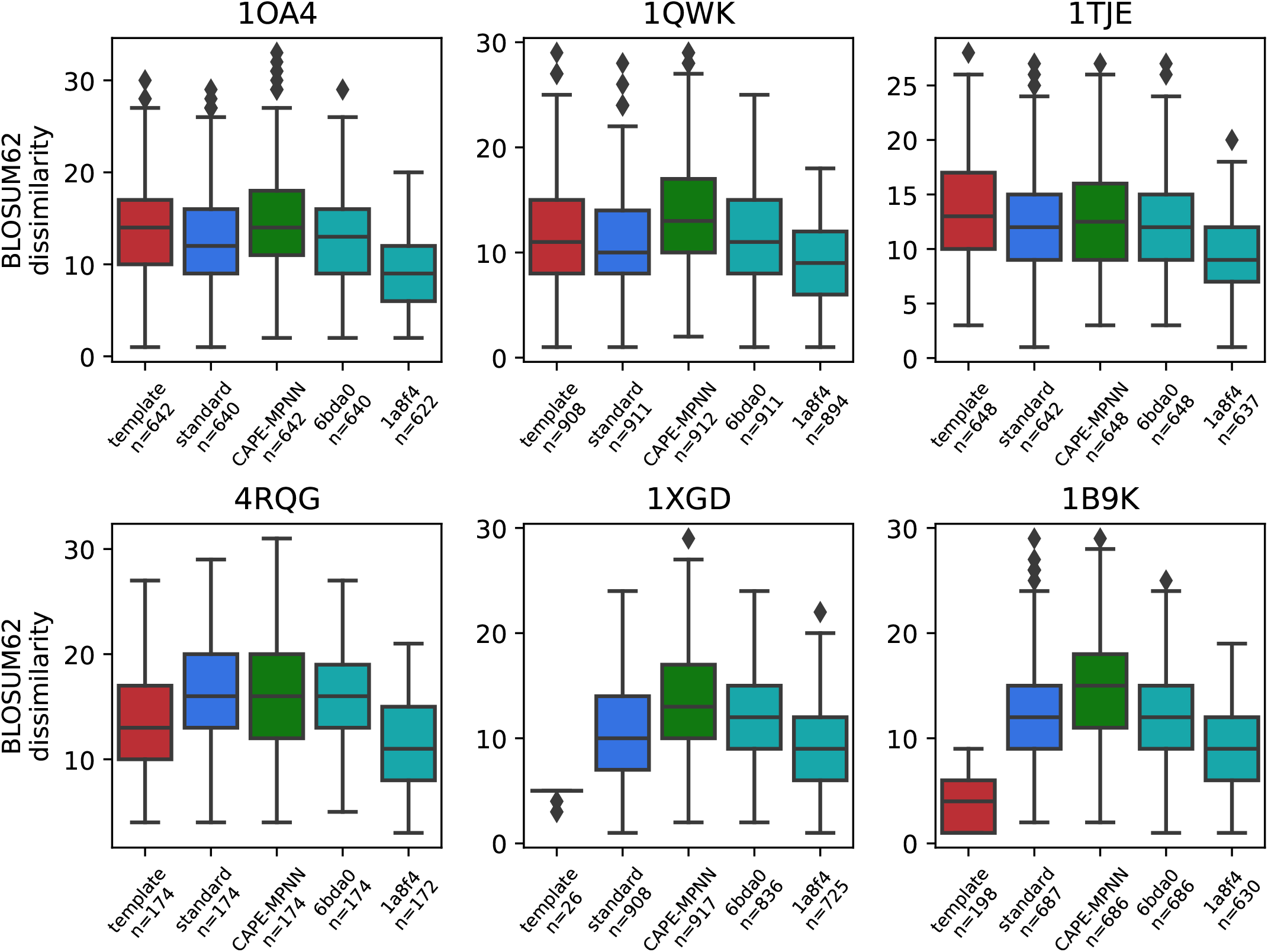
CAPE-Beam 6mers generated 8-10mers look most similar to the self-proteome for non-human proteins. Dissimilarity to the human proteome is predictive of immunogenicity [22]. The figures shows the BLOSUM62 dissimilarity (see **Section 4.4.2**) distribution of the non-self 8-10mers for the six validation set proteins. Five sequence sources are plotted: the original template protein, using standard (greedy) `ProteinMPNN` decoding, using `CAPE-MPNN` and using `CAPE-Beam` requiring either all 5mers or all 6mers (and shorter) to come from the human proteome. The number *n* below the label shows how many of the 8-10mers are non-self. We see that forcing all 6mers and shorter to come from the human proteome generated the least dissimilar 8-10mers for the bacterial (1OA4, 1TJE), nematoda (1QWK) and reptilia (4RQG) proteins. Only in the case of the *Homo sapiens* (1XGD) and *Mus musculus* (1B9K) proteins, the original template proteins are more similar. This is no surprise, since they are either strong homologues or actually from the human genome. However, `CAPE-Beam` 6mers always generated less dissimilar kmers than only forcing 5mers from the human proteome or the standard decoding approach.

Finally, **Figure 4** then focuses on some molecular properties of generated sequences. Based on the low TM scores observed in **Figure 2** for requiring all 7mers to come from the human proteome, we only consider `CAPE-Beam` designs with `min_self_kmer_length` of 5 and 6 here. We also only keep the `CAPE-Beam 0.5` designs. For the rosetta energy per AA, we find that these are (except for outliers) acceptable for the designs requiring only 5-mers from the proteome. Also, many of the designs that require 6-mers from the human proteome come have rosetta energy below −1. With regards to aggregation score, we find that the designs all are in a comparable area to the original proteins. There are some differences with regards to isoelectric point, which would need further investigation.

**Figure 4.**
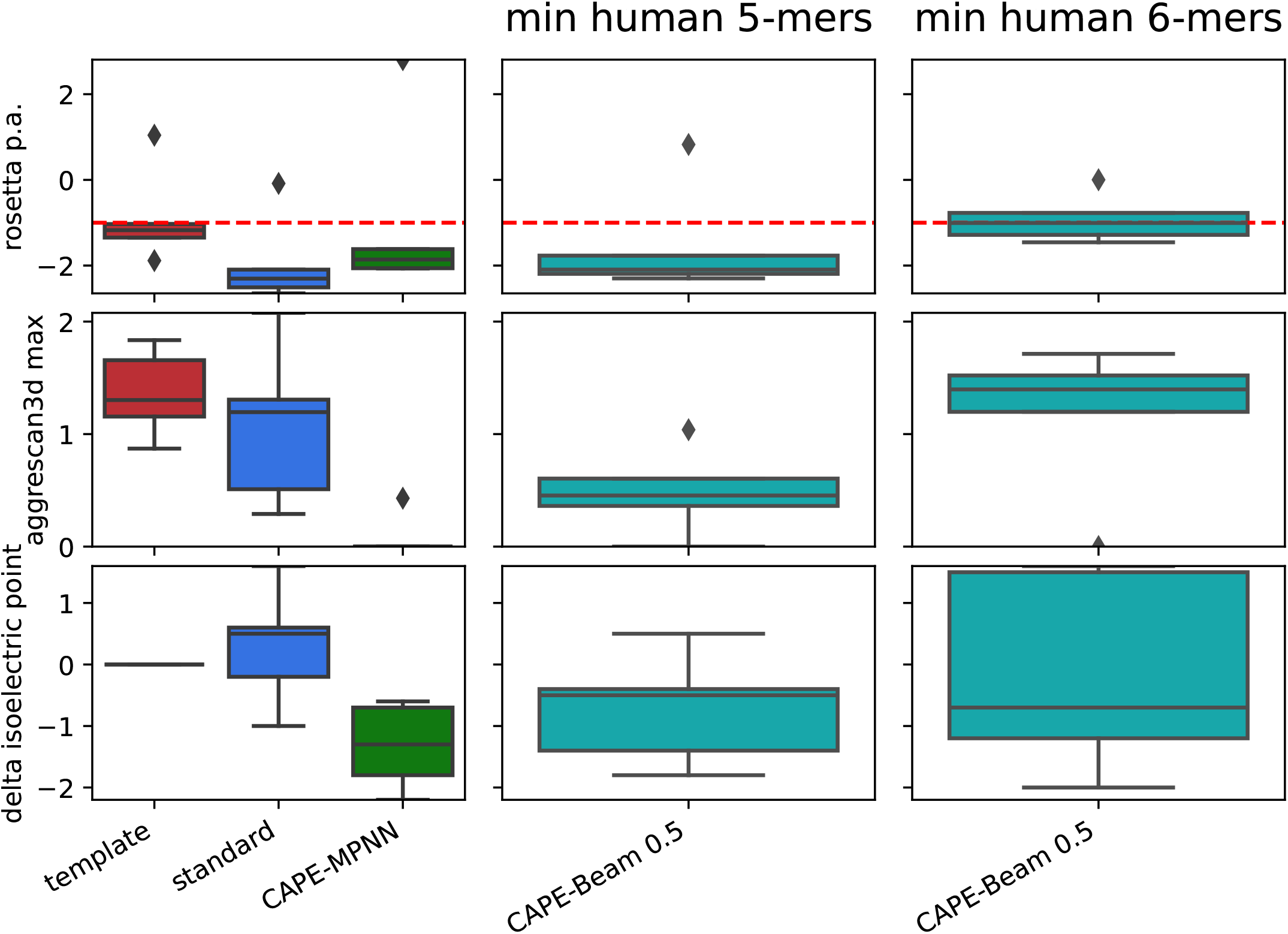
Molecular properties of generated sequences. For the validation set proteins, this figure compares the distribution of Rosetta scores per AA, aggregation scores and difference in iso-electric point (see **Section 4.4.3**) between the original template protein, using greedy (standard) ProteinMPNN decoding, using `CAPE-MPNN` and either forcing requiring only 5mers and shorter or also 6mers to come from the human proteome for CAPE-Beam decoding. With regards to stability (rosetta p.a.), wee see that `CAPE-Beam` 5mer shows similar values to `CAPE-MPNN` while only about half of the CAPE-Beam 6mer designs fall into the typical score range for a refined structure. The aggregation scores all seem to be in the range or better than the original templates and `CAPE-Beam` designs seem to be more similar to the original protein with regards to isoelectric point than `CAPE-MPNN` designs.

## 6 Discussion

In this work we have shown that using a modified sampling approach from `ProteinMPNN` allows us to generate proteins that only consist of kmers up to 10 that are either predicted not to be presented to CTLs or can be found in the human proteome and will, therefore, be subject to central tolerance. In addition, for some proteins we have shown that as an additional safety measure we can increase the length of kmers making up the generated proteins that come from the human proteome. Although we were unable to extend the length to cover all peptides that are presented via MHC-I, this should still be beneficial in light of results that show that TCRs bind far less strongly to tumor-associated antigenic peptides, which have very similar sequences to those peptides found in the human proteome, than to viral ones [5]. We have also shown that this reduces the “BLOSUM62 dissimilarity” from non-self kmers to the human proteome - which according to Richman et al. [22] tends to reduce their immunogenicity.

We have compared our approach to a fine-tuning approach for `ProteinMPNN` called `CAPE-MPNN` and find that `CAPE-Beam` has a higher potential to reduce immunogenicity in precision medicine due to its ability to force the predicted number of presented non-self 8-10mers to be zero. It also holds high potential to reduce immunogenicity in a broader population due to its ability to produce designs consisting of 8-10mers that are more similar to the proteome.

### Limitations

One of the limitations in this work is that we have only carried out the analysis for the genotype of one specific hypothetical patient. However, the alleles used are very common. Also, some immunogenic peptides might have longer length than the canonical 8-10 AAs. However, in many of these cases a 8-10 AA long core can be found which would be covered by our method. Future work will also have to include laboratory confirmation of the designs instead of relying solely on bioinformatics prediction methods.

## Acronyms

AA: amino acid
Ab: antibody
AR: auto-regressive
CTL: cytotoxic T-lymphocyte
GNN: graph neural network
LLM: large language model
MHC-II: MHC Class II
MHC-I: MHC Class I
ML: machine learning
MPNN: message passing neural network
pI: isoelectric point
PWM: position weight matrix
REU: Rosetta Energy Unit
SOTA: state of the art
T-cell: thymus dependent lymphocyte
TCR: T-cell receptor
VAE: Variational Autoencoder

## 7 Acknowledgements

This work was supported by the United Kingdom Research and Innovation (grant EP/S02431X/1), UKRI Centre for Doctoral Training in Biomedical AI at the University of Edinburgh, School of Informatics. For the purpose of open access, the author has applied a creative commons attribution (CC BY) licence to any author accepted manuscript version arising.

This project was supported by the Royal Academy of Engineering and the Office of the Chief Science Adviser for National Security under the UK Intelligence Community Postdoctoral Research Fellowship programme.

J.A. Alfaro was supported by (i) United Kingdom Research and Innovation (grant EP/S02431X/1) (UK-IC post-doctoral fellowship), (ii) the project ‘International Centre for Cancer Vaccine Science’ which is carried out within the International Agendas Program of the Foundation for Polish Science, cofinanced by the European Union under the European Regional Development Fund, and (iii) The JunG Grant, awarded to Dr. Javier Alfaro at the University of Gdansk. The authors would like to thank ‘CI-TASK, Gdansk’, and the ‘PL-Grid Infrastructure, Poland’ for providing their hardware and software resources. Dr. Rajan was supported by the KATY Consortium H2020-SCI-FA-DTS-2020-1.

## 8 Author contributions

**Hans-Christof Gasser:** Conceptualization, Methodology, Software, Validation, Writing - Original Draft, Writing - Review & Editing, Visualization **Ajitha Rajan:** Conceptualization, Writing - Review & Editing, Visualization, Supervision **Javier Alfaro:** Conceptualization, Writing - Review & Editing, Visualization, Supervision

## Footnotes

1 https://seq2fun.dcmb.med.umich.edu/TM-align/

